# Role of small non-coding RNAs in DSB repair in *Arabidopsis thaliana*

**DOI:** 10.64898/2026.10.09.757836

**Authors:** Etienne Mochet, Charles White, Jérémy Verbeke, Heïdi Serra, Olivier Da Ines, Sébastien Lageix

## Abstract

DNA double-strand breaks (DSBs) are toxic lesions that must be repaired to avoid genetic loss and genome rearrangements. Various conserved pathways are involved in DSB repair, producing different outcomes. Accumulating evidence has identified RNA, particularly small RNAs as a new player in DSB repair. DSB-induced small RNAs (diRNAs) have been detected at DSB-flanking sequences using reporter systems, but their endogenous occurrence and function in DNA repair remain debated. Here, we investigated diRNA biogenesis in *Arabidopsis thaliana* using CRISPR/Cas12a to induce DSBs at three transcribed loci and one untranscribed locus. We found that 21-nt diRNAs are generated at transcribed loci, in a Dicer-like 4 (DCL4)-dependent manner, but not at the untranscribed locus, indicating that active transcription is required for diRNA production. Long-read DNA sequencing analysis showed that diRNAs do not detectably influence DSB repair via either end-joining or microhomology-mediated end-joining. Furthermore, endogenous diRNAs have a modest effect on homologous recombination-associated repair, as impairment of the diRNA pathway in *ago2* mutants caused only a slight reduction in RAD51 focus formation following Cas12a-induced DSBs. Together, these findings indicate that diRNAs are not essential for DSB repair but may function in post-transcriptional silencing of damaged genes, potentially preventing aberrant transcription until repair is completed.

## Introduction

The DNA of all living cells is continuously threatened by damage arising from both exogenous environmental stressors and endogenous metabolic processes. Efficient repair of these lesions is essential to preserve genome integrity and ensure cellular viability. Among the various types of DNA damage, double-strand breaks (DSBs) are particularly cytotoxic because they involve the simultaneous cleavage of both DNA strands, leading to potential loss of genetic information and precluding simple repair using an intact complementary strand. If unrepaired or misrepaired, DSBs can cause chromosome rearrangements, deletions, or loss of chromosomal segments, thereby compromising genomic stability (1).

To cope with these threats, organisms have thus evolved multiple sophisticated DNA repair pathways (2, 3). DSBs are primarily repaired through two major pathways: non-homologous end joining (NHEJ) and homologous recombination (HR). NHEJ directly ligates broken DNA ends and is often associated with small insertions or deletions (2, 4). In contrast, HR is a more accurate process that uses an intact homologous DNA molecule as a template to restore the damaged molecule (2, 5). Central to HR is the generation of single-stranded DNA (ssDNA) by end resection, followed by binding of Replication Protein A (RPA), and subsequent replacement of RPA by the RAD51 recombinase. This process leads to the formation a nucleoprotein filament that catalyzes homology search and strand invasion of a homologous DNA molecule that will be used as a template to repair the break (2, 5).

Remarkably, accumulating evidence over the past decade has revealed an unexpected association between RNA molecules, particularly small RNAs, and DSB repair (6–15). The first evidence came from a study in the filamentous fungus *Neurospora crassa*, in which DNA damage was shown to induce the production of QDE-2–interacting RNAs (named qiRNAs) derived from ribosomal DNA repeats, defining a novel class of small interfering RNAs associated with genome stability (6). Subsequently, DSB-induced small RNAs (diRNAs) were detected in both *Arabidopsis thalian*a and human cells. In Arabidopsis, diRNAs, predominantly 21-nucleotide siRNAs, were identified as being produced at sequences flanking a site-specific DSB introduced into a transgene (15). Similar small RNAs were detected at DSB sites in human cells, implying an evolutionarily conserved phenomenon (7, 15). Finally, in Drosophila, small RNAs complementary to DNA damage sites were shown to be required for a proper DNA Damage Response (DDR), while the introduction of synthetic small RNAs matching the break site was sufficient to rescue DDR signaling, further supporting a functional role for small RNAs in DNA damage repair (8).

Notably, diRNAs are produced from both sense and antisense strands surrounding the break site at similar frequencies, suggesting their origin from double-stranded RNA precursors, potentially generated by RNA-dependent RNA polymerases. In Arabidopsis, diRNA biogenesis requires the RNA polymerases IV and V, the ATR kinase, and Dicer-like (DCL) enzymes. Accordingly, loss of these components, as well as ARGONAUTE 2 (AGO2), impairs homologous recombination efficiency, suggesting that diRNAs participate in DSB repair (15–17).

Mechanistic insight into how diRNAs influence DSB repair later emerged from the identification of downstream RNA-binding factors. The double-stranded RNA-binding protein INVOLVED IN DE NOVO 2 (IDN2), previously characterized for its role in RNA-directed DNA methylation (RdDM), physically interacts with both AGO2 and the replication protein A (RPA) complex. In *idn2* and *ago2* mutants, RPA abnormally persists at resected DNA ends, RAD51 loading is compromised, and HR is defective. These observations support a model in which AGO2–diRNA complexes recruit IDN2 to DSBs, facilitating displacement of RPA from single-stranded DNA and promoting RAD51 filament formation (18). More recently, the conserved SMC5/6 complex, essential for genome stability, was shown to be recruited to DSBs by a protein scaffold composed of IDN2, the DNA damage response factor CDC5, and the transcriptional coactivator ADA2b. Assembly of this scaffold requires diRNAs generated from sequences surrounding the break, as well as functional AGO2 and the RNA-binding activity of IDN2, indicating that small RNAs can nucleate multiprotein repair complexes at DSBs. Altogether, these studies support a model in which 21-nt small diRNAs recruit and/or stabilize DNA repair factors at DSB sites likely through sequence complementarity (16).

However, robust production of 21-nt small diRNAs has so far been convincingly demonstrated only at transgenic reporter loci or repetitive sequences, most commonly using GUS-based reporter constructs (15, 18). Consequently, whether similar 21-nt small diRNAs are robustly generated at endogenous genes following DSB formation remains unclear. Indeed, analysis of CRISPR/Cas9 or TALEN-induced DSBs in Arabidopsis and rice confirmed 21-nt diRNA production at a transgene break site but failed to detect comparable small RNAs at endogenous loci (19). Notably, diRNA production at transgenes required active RNA polymerase II transcription at the damaged locus and depended on canonical post-transcriptional gene silencing factors Dicer-like 4 (DCL4) and RNA-dependent RNA polymerase 6 (RDR6) but not Dicer-like 2 (DLC2). Mutants defective in these factors did not show defects in DSB repair, leading to the conclusion that 21-nt small diRNAs are not strictly required for repair under these conditions (19). Recently, a genome-wide study using an inducible SbfI restriction enzyme system detected widespread transcription initiation from DSBs across the Arabidopsis genome, particularly within actively transcribed genes (20). These damage-induced RNAs were often bidirectional and produced by RNA polymerase II. However, although referred to as diRNAs, these molecules were identified by poly(A)+ RNA sequencing and therefore represent long transcripts rather than the 21-nt small diRNAs originally described as diRNAs. This distinction is important as long DSB-induced RNAs are Polymerase II-dependent while small diRNAs production is mediated by Pol IV/V (15). These observations raise the intriguing possibility that such long RNAs produced by Polymerase II could serve as templates for the generation of a precursor double-stranded RNA by RDR6, that will eventually be processed by DCL4 into 21-nt small RNAs (19).

Small RNAs (21-nt) have also been implicated in the repair of UV-induced DNA lesions in plants (21). Following UV-C exposure, Arabidopsis produces 21-nt small RNAs at many sites of pyrimidine dimers in a Pol IV–RDR2–DCL4-dependent manner. These UV-induced siRNAs associate with ARGONAUTE1 (AGO1) and the damage recognition factor DDB2, suggesting a functional role for 21-nt small RNAs in nucleotide excision repair. However, UV-induced photoproducts are chemically and molecularly distinct from DSBs, and thus these findings do not directly address 21-nt small diRNAs production and function at DSBs.

Collectively, existing studies support the hypothesis that 21-nt small diRNAs can be produced in response to DNA damage at transgenes and may influence repair processes in plants. However, whether such 21-nt small diRNAs are generated at endogenous DSBs, and whether they directly promote recombinational repair at native genomic loci, remains an open question. In particular, although a role for diRNAs in HR has been suggested, their potential involvement in NHEJ has not yet been directly tested in plants.

Here, we used CRISPR/Cas12a to induce targeted DSBs at three actively transcribed loci and one untranscribed locus in *Arabidopsis thaliana* to characterize diRNA production and test their functional role in DSB repair. We find that 21-nt diRNAs are generated at transcribed loci but not at the untranscribed region, indicating that active transcription is required for diRNA biogenesis. We further show that diRNAs are produced in a Dicer-like 4 (DCL4)-dependent manner and that, consistent with their known role in gene silencing, the presence of diRNAs appears to be associated with post-transcriptional gene silencing (PTGS) of damaged genes. Remarkably, long-read DNA sequencing analyses show that these diRNAs do not detectably influence DSB repair via either end-joining or microhomology-mediated end-joining. Furthermore, our data indicate that endogenous diRNAs have only a minor effect on the HR-associated response to DSBs, as impairment of the diRNA pathway in *ago2* mutants resulted in only a slight reduction in RAD51 focus formation following Cas12a-induced DSBs. Together, our results suggest that diRNAs are not essential for DSB repair *per se,* but may instead function in the silencing of damaged genes via PTGS, thereby preventing aberrant transcript accumulation at damaged genes until proper repair has occurred.

## Materials and Methods

### Plant material and growth conditions

All *Arabidopsis thaliana* plants used in this study were in the Columbia (Col-0) ecotype background. The following mutant lines were used: *dcl4-2* (GABI-Kat, GK-160G05) (65) and *ago2-1* (SALK_037548) (15).

For culture, seeds were stratified in water at 4 °C for 2 days and grown either on soil or *in vitro* in a growth chamber. For *in vitro* culture, seeds were surface sterilized for 5 min with 75% Ethanol, 0.05% SDS, rinsed with 95% Ethanol for 5 min and air-dried. Sterilized seeds were then sown on Murashige and Skoog (MS) medium, stratified at 4 °C for 2 days and placed in a growth cabinet. All plants were grown under 16h light /8 h dark cycles at 23 °C and 60% relative humidity. For DSB induction, plants were cultivated for 10 days at 29°C under the same 16/8h light/dark cycle, and 60% relative humidity.

### Generation of FnCas12a-expressing plants

To induce DSBs, we used our CRISPR/Cas12a system as previously described (22). *Arabidopsis thaliana* Col-0 plants stably transformed with the FnCas12a T-DNA construct were crossed with *dcl4* and *ago2* mutant plants, and analyses were performed on their F2 or F3 progenies.

As a negative control in which DSB cannot be induced (referred to as “+Cas, -gRNA”), Col-0 plants were also transformed with a construct expressing only the FnCas12a endonuclease (66). Additionally, FnCas12a constructs were digested with ScaI and StuI restriction enzymes to truncate 790 amino acids of the Cas12a protein (from 11^th^ to the 800^th^ amino acid). The resulting truncated FnCas12a T-DNA construct *(ΔCas*) was transformed into *Arabidopsis thaliana* using the floral dip method (67) and served as an additional negative control.

### Genomic target loci, T-DNA construction and plant transformation

In addition to our two previously described targeted genomic loci: LPP2 (At1G5080) and Chr3_92 (located in the repeat-rich, pericentromeric region of Chromosome 3 (Chr3:12113075-12114245), and consisting of two perfect 142 bp tandem direct repeats separated by 742 nt (Fig. 1; (22)), we used two additional targeted loci: *XPF* (At5G41150) and *ERCC1* (At3G05210).

**Figure 1.**
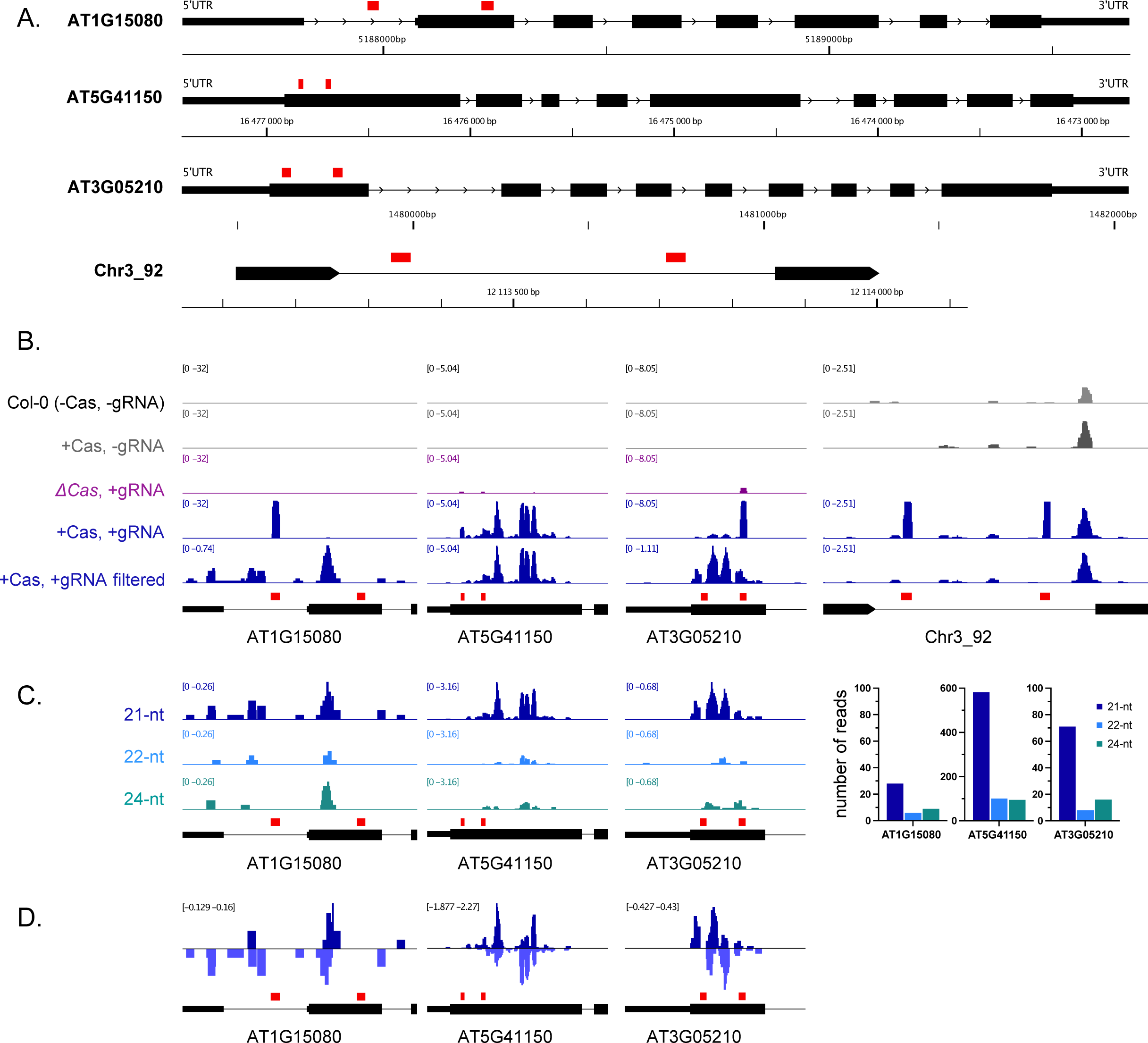
DSBs at Transcribed Euchromatic loci but not Non-Transcribed Heterochromatic region trigger 21-nt Small RNA Production. (A) Schematic representation of the 3 expressed genes and the pericentromeric repeat-rich region (“Chr3_92”) targeted by the CRISPR/Cas12 system. The two CRISPR/Cas12 cut-sites are indicated by red boxes. (B-C-D) Integrated Genome View (IGV) of small RNA sequencing tracks at the CRISPR/Cas12-targeted regions (counts per million, CPM). *AT1G15080*, *AT5G41150*, *AT3G05210,* and *Chr3_92* (B) Accumulation of small RNAs at the chosen targets in different genetic backgrounds. Small RNAs perfectly matching the CRISPR/Cas12 cut-sites were filtered out as they could not be distinguished from CRISPR guide RNAs and to allow a better visual representation of the small RNAs after DSB induction. For Chr3_92 target, the ΔCas +gRNA line was not generated, and therefore the data are not available. (C) Different populations of diRNAs of 21 nt, 22 nt and 24 nt at endogenous loci. Bar-graph shows the number of reads mapping to a window containing the two CRISPR/Cas12 cut-sites, regions where diRNAs are exclusively observed. (D) Positive values represent sense 21-nt diRNAs while negatives are antisense 21-nt diRNAs at the targeted exons.

For each target, two gRNA constructs were designed to induce DSBs simultaneously in both target genes. The gRNA were cloned into the expression plasmids kindly provided by Dr. Seiichi Toki’s group, following previously described methods (66). The gRNAs were selected to induce two distinct DSBs in the first exon of both targets (Fig. 1). Their sequences are provided in Supplementary Table S1. *Arabidopsis thaliana* were transformed by the floral dip method (67).

### RNA isolation

Total RNA was extracted from a pool of ten 10-day-old plantlets using RNAzol RT (RN190, Molecular Research Center). RNA concentration and quality were assessed using the Qubit fluorometer (RNA High Sensitivity kit, Thermo Fisher) and a Nanodrop spectrometer, respectively.

Both Small RNA-seq and RNA-seq libraries were generated from at least 1μg of total RNA. UMI Small RNAseq (SE50) and Transcriptome Strand-specific sequencing (PE150) were performed on a DNBSEQ platform at Beijing Genomics Institute (BGI, Hong Kong).

### Small RNA-seq analysis

Adapters trimming and sequence quality check was performed by the BGI platform, using their SOAPnuke(68) software with the following parameters: -n 0.01 -l 20 -q 0.1 --highA 1 --minReadLen 15 --maxReadLen 44 --ada_trim. Trimmed reads were aligned to the *Arabidopsis thaliana* reference genome (TAIR10; https://www.arabidopsis.org/) using STAR (69) (2.7.11b) with the following parameters: STAR --outFilterMatchNmin 10 --alignIntronMa× 10000 – alignSJoverhangMin 3 --outFilterMismatchNmax 0. To rule out possible reads mapping to the FnCas12a T-DNA construct, only reads with a perfect CIGAR score were kept using SAMtools (70) (1.22.1) and reads perfectly mapping to the CRISPR-Cas recognition sites were removed using BEDTools (71) intersect (v2.21.1). Size selection (21, 22 or 24-nt) and sense analysis were performed on these processed BAM files using SAMtools. For an accurate representation of the different small RNA populations (21-nt, 22-nt and 24-nt) in Integrative Genomics Viewer (IGV (72)), a scaling factor (1/#mapped reads)*10^6 was applied when using BamCoverage (73).

21-nt reads, from F3 and T1 plants, spanning the diRNAs generation sites were combined and reverse reads were reverse-complemented, so all the reads are in the same orientation. Nucleotide frequency was assessed at both position 1 (5′ bias) and 21 (3′ bias).

### RNA-seq analysis

As above, read trimming and quality check was done by BGI with the following parameters (-n 0.01 -l 20 -q 0.4 --adaMR 0.25 --polyX 50 --minReadLen150) and trimmed reads were aligned to TAIR10 with STAR (69) (2.7.11b, --outFilterMatchNmin 10 --alignIntronMa× 10000 – alignSJoverhangMin 3). BamCoverage (deepTools) was used to generate CPM-normalized BigWig coverage files, which were subsequently visualized using IGV. FeatureCounts was used to count paired end reads for each target gene with the Araport11 genome annotation. Transcripts Per Million (TPM) were calculated with the following formulas:

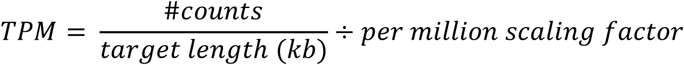

The per million scaling factor was obtained by diving the sum of all reads per kilobase (RPK) in a sample by 10^6^.

### RTqPCR

Two μg of total RNA were treated with RQ1 RNAse-Free DNase (Promega) and reverse-transcribed using M-MLV Reverse Transcriptase (Promega) with both random primers and oligo-dTs.

Real-time RT-qPCR amplification was carried out with 2 μL of cDNA using TB Green Premix Ex Taq II (Tli RNaseH Plus, TaKaRa) in a total volume of 20 μL using a LightCycler 480 system (Roche), following the manufacturer’s recommendations. Data were normalized to the *ACTIN2* (At3G18780) reference gene and relative expression of each target was assessed with the 2 ^-ΔCt^ method from four independent biological samples.

### Nanopore DNA sequencing

Genomic DNA was extracted from individual 10-day-old plantlets, using the Promega Wizard Genomic DNA Purification Kit and double-stranded DNA concentration was assessed by the Qubit fluorometer (High Sensitivity kit, Thermo Fisher). The CRISPR-Cas12a cutting efficiency was evaluated by PCR amplification of the region spanning the two cut-sites for each target loci (22).

Libraries for Nanopore sequencing were generated as described in our previous work (22). Briefly, 100 ng of pooled genomic DNA from five plantlets were amplified by PCR using Platinum SuperFi II DNA Polymerase (Invitrogen) and purified using Agencourt AMPure XP beads. Libraries were prepared for Nanopore sequencing using the SQK-NBD 114 ligation sequencing kit according to the manufacturer’s protocol and sequencing was performed with a MinION Mk1c device using R10.4.1 (FLO-MIN 114) flow cells. The Nanopore data were basecalled using the SUP parameter of Dorado v.0.7.0 (https://github.com/nanoporetech/dorado).

NHEJ and MMEJ analysis were performed using an in-house Python and R script as published previously (22) and is publicly available at https://github.com/chrlesw/dsbrepair. Reads with a deletion ≤ 2 bp were ruled out from the analysis as they could not be distinguished from Nanopore sequencing errors.

### Immunolocalization on root nuclei

For immunolocalization of RAD51, seedlings were grown for 10 days at 23°C on MS medium, then incubated at 29°C for 4 hours to induce DSBs. For MMC induction, 5-day-old seedlings grown on half-strength MS medium were transferred on half-strength MS liquid medium supplemented with 30 μM MMC and further grown for 6 hours. Samples were fixed in 4% PFA-1X PME buffer for 45 min. Immunostaining in root tip nuclei was then performed as previously described (74). Slides were incubated with rat α-RAD51 (diluted 1/500 in 3% BSA, 0.05% Tween-20 in 1X PBS) in a moist chamber at 4°C overnight. Slides were washed three times in 1X PBS-0.05% Tween-20, airdried, and then incubated with secondary antibody solution diluted 1/1,000 in 3% BSA, 0.05% Tween-20 in 1X PBS in a moist chamber for 3 h at RT in the dark. Slides were finally washed three times in 1X PBS0.05% Tween-20, air-dried, and mounted in VECTASHIELD mounting medium containing DAPI (1.5 μg/ml; Vector Laboratories Inc.).

All observations were made with a motorized Zeiss AxioImager.Z1 epifluorescence microscope (Carl Zeiss AG, Germany) using a PL Apochromat 100X/1.40 oil objective. Photographs were taken with an AxioCam Mrm camera (Carl Zeiss AG, Germany) and appropriate Zeiss filter sets adapted for the fluorochromes used: filter set 25HE (DAPI), filter set 38HE (Alexa 488), and filter set 43HE (Alexa 596). Image Z-stacks were captured in three dimensions (x, y, z) and further deconvoluted with the deconvolution module (theoretical PSF, iterative algorithm) of the Zeiss ZEN lite software (Carl Zeiss AG, Germany). For presentation, pictures are collapsed Z-stack projections obtained using the Extended-focus module (projection method) of the Zeiss ZEN lite software.

## Results

### DSBs in Transcribed Euchromatin loci but not Non-Transcribed Heterochromatin region trigger 21-nt Small RNA Production

To investigate the production of diRNAs at endogenous loci, we used the CRISPR/Cas12a system to induce 5′-overhang DSBs at transcribed and non-transcribed loci in the *Arabidopsis thaliana* genome (Fig. 1A). We targeted the euchromatic, transcriptionally active gene *LPP2* (AT1G15080), located in the central region of the left arm of chromosome 1. To do so, two guide RNAs (gRNAs) were designed to induce DSBs within the first intron and the second exon, respectively. In addition to this transcribed locus, a non-transcribed pericentromeric locus (labelled Chr3_92) was also targeted (Fig. 1A). This target locus lies in the repeat-rich, pericentromeric region of chromosome 3 and consists of two perfect 142 bp tandem direct repeats separated by 742 nt. Two unique Cas12a target sites were chosen in the sequences between the two repeats, such that the induced DSBs are flanked by tandem direct repeat DNA sequences. We have previously shown that the majority of repaired DSBs at these two targets carried small deletions, consistent with local repair through end-joining mechanisms (22).

To assess diRNA production, we performed small RNA sequencing on plants in which Cas12a-induced DSBs were generated either at the euchromatic, transcriptionally active locus or at the non-transcribed heterochromatic target. Because Cas12a is active at 29°C, all seedlings were grown for 10 days at this temperature to promote robust DSB induction (see Methods). In the absence of DSB induction (-Cas -gRNA and +Cas -gRNA), no small RNAs were detected at the transcribed *LPP2* locus (AT1G15080, Fig. 1B). In contrast, low levels of small RNAs were detected at the heterochromatic target, consistent with their natural and constitutive presence in this genomic context.

In plants expressing the guide RNAs (gRNAs) and a truncated, non-functional Cas12 *(ΔCas12,* +gRNA), we observed an accumulation of small RNAs at the gRNA target recognition sites (AT1G15080, Fig. 1B). These small RNAs do not correspond to diRNAs, as no DSBs are generated without Cas12 activity. Instead, they represent by-products of gRNA expression and/or small RNAs produced in response to the T-DNA, which encodes the gRNAs among other elements. Consistent with this hypothesis, these small RNAs also showed perfect sequence matches to the T-DNA. Consequently, for the remainder of this study, small RNAs mapping to gRNA recognition sites were filtered out and excluded from subsequent analyses. Remarkably, following DSB induction in CRISPR/Cas12a expressing plants (+Cas, +gRNA), a clear accumulation of diRNAs was observed exclusively at the *LPP2* target locus (AT1G15080, Fig. 1B). Notably, diRNA accumulation was enriched in both exonic and intronic regions surrounding the Cas12a cleavage sites. Classification of the diRNAs according to their size indicates that these diRNAs predominantly correspond to 21-nt species (Fig. 1C) and were generated from both sense and antisense strands (Fig. 1D). In contrast, apart from gRNA by-products, no detectable small RNA enrichment was detected at the heterochromatic, non-transcribed Chr3_92 target following DSB induction (+Cas, +gRNA) (Fig. 1B). We note that consistently with the heterochromatic context, this locus naturally produces siRNAs, predominantly 24-nt species. However, their accumulation was not DSB-dependent, as these siRNAs were also detected in the absence of DSB induction (Fig. 1B).

These results suggest that diRNA production occurs specifically at endogenous transcribed loci. To confirm this, we performed small RNA sequencing on plants in which Cas12a-induced DSBs were induced at two additional euchromatic, transcribed loci: AT5G41150 (*AtXPF)* and AT3G05210 (*AtERCC1)*. For each gene, two gRNAs were designed to introduce DSBs within the first exon (Fig. 1A). In both cases, strong accumulation of diRNAs was detected at the target loci. These diRNAs were detected exclusively within exons harboring Cas12a-induced DSB, predominantly consisted of 21-nt species, and originated from both sense and antisense strands (Fig. 1B-D).

Collectively, these results establish, to our knowledge, the first evidence of robust 21-nt diRNA production from endogenous transcribed genes in *Arabidopsis thaliana*, demonstrated across three independent target loci, while revealing a striking dependence on transcription, as no diRNAs were detected at an untranscribed, heterochromatic locus.

### diRNAs production requires the essential PTGS factor DCL4

To assess the biogenesis pathway of these diRNAs, we performed small RNA sequencing analysis in a *dcl4* mutant background. DCL4, a key component of the RNA silencing machinery responsible for processing double-stranded RNA into 21-nt small RNAs, has previously been shown to be required for the production of diRNAs targeting a reporter transgene in *Arabidopsis thaliana* (15, 19). Consistent with this, absence of DCL4 leads to strong reduction in diRNA accumulation at all three euchromatic, transcribed targets, demonstrating that the DSB-induced small RNA production at endogenous gene loci is DCL4-dependent (Fig. 2A).

**Figure 2.**
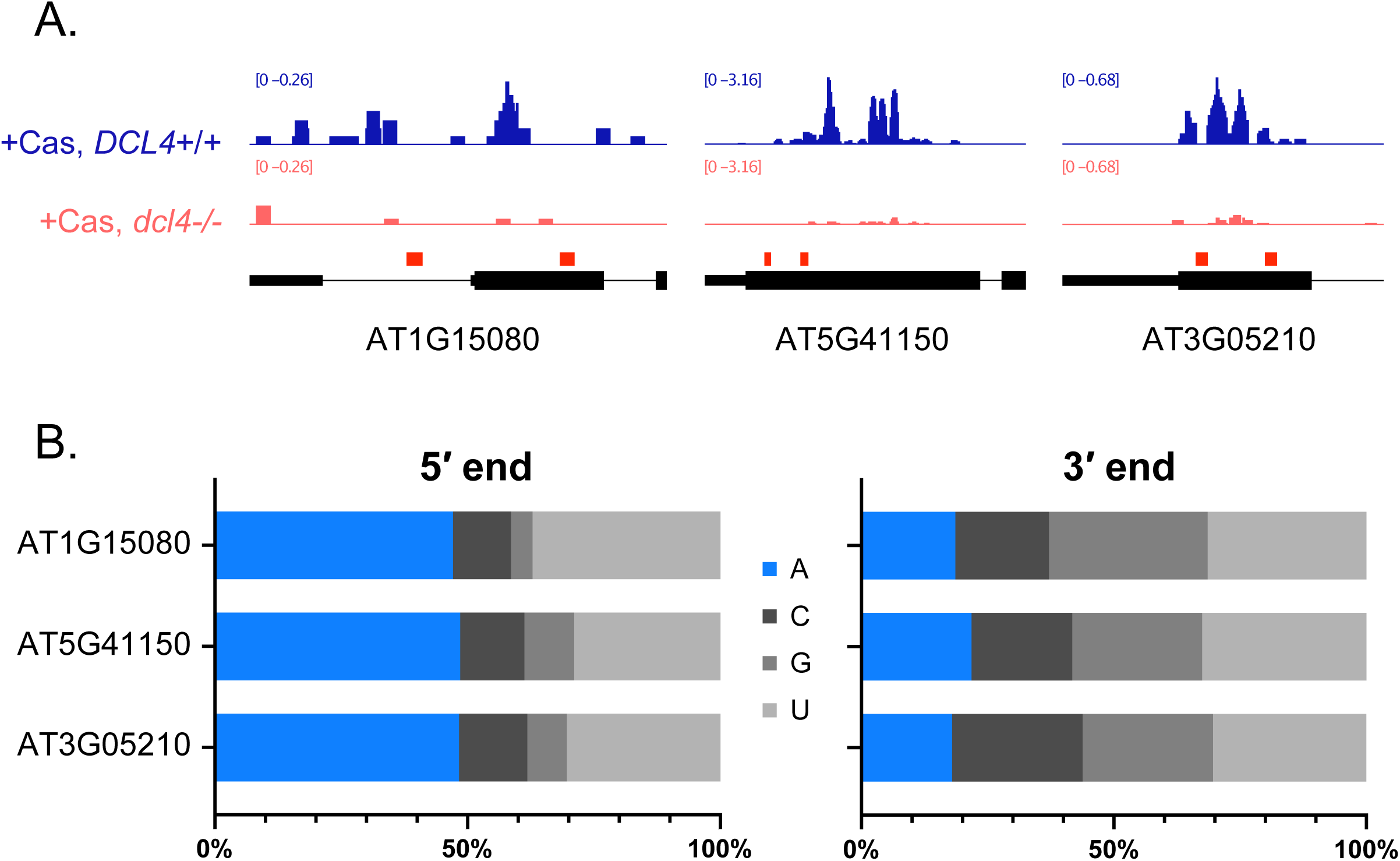
diRNA biogenesis is DCL4-dependent, and their sequence shows signature of AGO2 loading. (A) Integrated Genome Viewer of 21-nt diRNAs at the three CRISPR/Cas12-targeted genes in *DCL4* wild-type and mutant (*dcl4*−/−) backgrounds. CRISPR/Cas12 cut-sites are represented by red boxes. (B) Nucleotide frequencies at the 5′ and 3′ ends of the 21-nt diRNAs, observed at the three CRISPR/Cas12-targeted loci, suggest a loading in AGO2. A= Adenine, C= Cytosine, G= Guanine, U= Uracil. AT1G15080 n=70 reads; AT5G41150 n= 913 reads; AT3G05210 n=89 reads.

Having established that these endogenous diRNAs are generated through a DCL4-dependent pathway, we next sought to determine their potential effector AGO proteins. Small RNAs associate with ARGONAUTE proteins to form effector complexes known as RNA-induced silencing complex (RISC), which are guided by sRNAs to complementary RNA targets. In *Arabidopsis thaliana*, ten AGO proteins have been identified, exhibiting distinct and selective sRNA loading preferences (23–28). This sorting specificity is largely governed by structural features of the AGO protein, notably the nucleotide specificity loop within the middle domain, which recognizes the 5′ nucleotide of the sRNA, thereby influencing AGO–sRNA binding preferences (29). Consistent with this, AGO1 preferentially associates with microRNAs bearing a 5′ uridine, AGO2 favors 21-nt RNAs with a 5′ adenine, AGO5 bound all three known size classes (21, 22, and 24 nt) of small RNAs, with bias toward 5′ cytosine and AGO4 predominantly binds heterochromatic 24-nt siRNAs with a 5′ adenine (23, 24). To identify the potential AGO partner of the 21-nt diRNAs characterized here, we analyzed their sequence composition, focusing on nucleotide frequencies at both termini (Fig. 2B). While no significant bias was observed at the 3′ end, we detected a pronounced enrichment for adenine at the 5′ end, with approximately 50% of sequences harboring a 5′ A. This strong bias suggests preferential loading of these euchromatic diRNAs into AGO2, consistent with previous findings (15).

### diRNAs negatively regulate the expression of damaged genes

Small RNAs are well known for their roles in transcriptional and post-transcriptional gene silencing (30–33). Given the accumulation of 21-nt small RNAs at DSB sites in transcribed genes, we hypothesized that these diRNAs could contribute to transcriptional repression and/or trigger post-transcriptional gene silencing (PTGS) at damaged loci. To test this, we analyzed our mRNA-seq data to assess the impact of diRNAs on transcript levels from targeted genes. In wild-type plants, where both DSBs and diRNAs are present following DSB induction, we observed a clear tendency towards reduced transcript accumulation compared with wild-type plants in which no DSBs were induced (Supplementary Fig. S1). In contrast, in a *dcl4* mutant background, where DSBs occur in the absence of diRNAs, transcript accumulation after DSB induction were comparable to those observed in no-DSB conditions. As this represents only one replicate, we sought to further validate these results by RT-qPCR using primers located downstream of the DSB sites (Fig. 3). Consistent with the mRNA-seq data, transcript levels were significantly reduced for two of the three target genes in wild-type plants when both DSBs and diRNAs were present compared with the corresponding wild-type plants in the absence of induced DSBs, while the third gene showed a similar but non-significant trend. Strikingly, this reduction was not observed in the *dcl4* mutant background, where DSB induction occurred in the absence of 21-nt diRNAs. Collectively, these results suggest that DSB-induced 21-nt diRNAs contribute to the reduction of transcript levels at damaged genes. Given the established role of 21-nt siRNAs in AGO-mediated post-transcriptional gene silencing, we hypothesize that these diRNAs may promote the post-transcriptional degradation of transcripts arising from DSB-containing loci, consistent with a PTGS mechanism rather than transcriptional repression.

**Figure 3.**
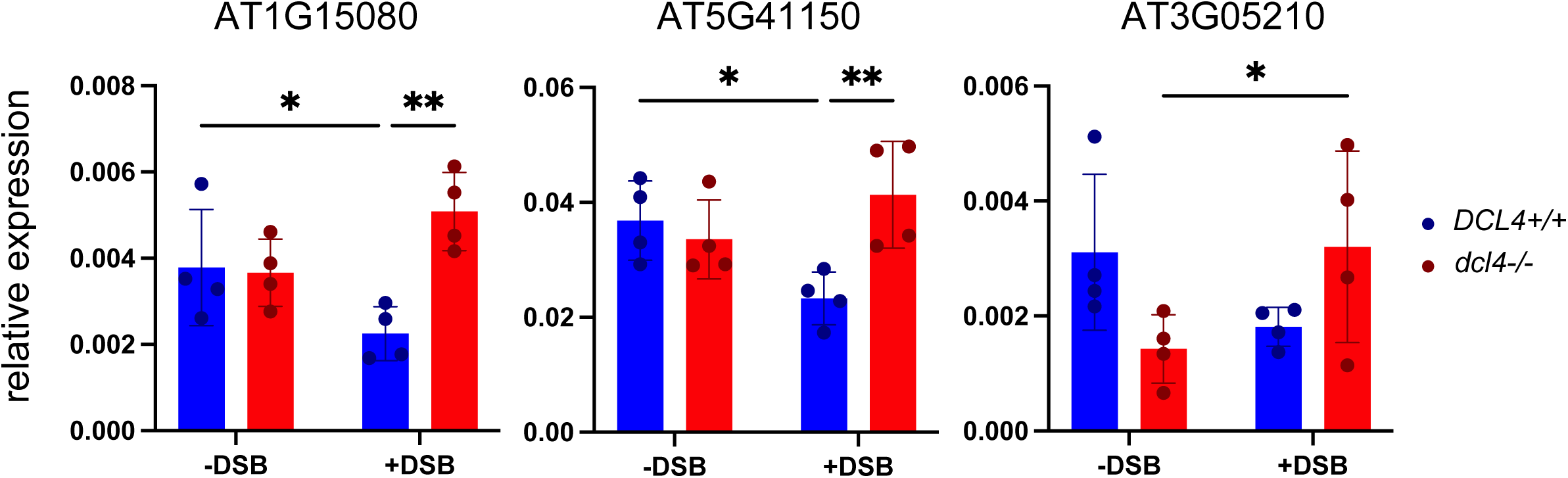
diRNA presence induces PTGS at damaged genes. The relative expression of CRISPR/Cas12 targeted genes was assessed by RTqPCR. Data were normalized to the *ACTIN2* reference gene, and the relative expression of each target gene to *ACTIN2* is presented (see methods). The “+DSB” annotation refers to lines expressing a functional CRISPR/Cas12 system (+Cas) targeting the evaluated gene, while “-DSB” is related to uninduced lines. Data were mean ± *s.d.*, n = 4 biological repeats. A two-way ANOVA with multiple comparisons was performed. \**p*-value < 0,05; \*\**p*-value < 0,001. Absence of annotation indicates no significant difference.

### diRNAs Have Little to No Impact on NHEJ and MMEJ

Beyond their role in transcription regulation, several studies have suggested that diRNAs contribute to DNA repair, particularly through homologous recombination (15, 16, 18, 34, 35). However, non-homologous end joining (NHEJ) is the predominant DSB repair pathway in many organisms, and the relationship between NHEJ and diRNA-mediated DNA repair remains unclear. To assess the potential role of diRNAs in NHEJ, we analyzed repair outcomes in 10-day-old seedlings at our eight Cas12a-induced DSB sites corresponding to the three euchromatic diRNA-producing loci and one non-transcribed locus, using Nanopore DNA sequencing. Consistent with our earlier observations, the majority of mutations induced by Cas12a at these loci were deletions, predominantly short in length (22). To quantitatively characterize these repair outcomes and avoid potential bias due to variation in cutting efficiency, deletion size distributions were calculated as the fraction of total deletions for each target site (Fig. 4 and Supplementary Fig. S2). In wild-type plants, deletion size distributions varied markedly between genomic loci, as well as between distinct Cas12a cleavage sites within the same locus, indicating that DSB repair outcomes are strongly influenced by local sequence context (Fig. 4 and Supplementary Fig. S2). We next compared deletion size distributions between wild-type (DCL4) and *dcl4* mutant (*dcl4*) plants to determine whether diRNAs influence NHEJ. The distributions were highly similar between the two genotypes, indicating that loss of diRNAs does not substantially alter the pattern or extent of deletions produced by NHEJ (Fig. 4A and Supplementary Fig. S2). AGO2 has previously been implicated in diRNA-mediated targeting to DSB sites and shown to associate with repair factors (15, 34). We therefore performed a similar analysis in *ago2* mutant and found no significant differences in the NHEJ profiles (Fig. 4B, Supplementary Fig. S2). Collectively, these results indicate that, although diRNAs are robustly produced at euchromatic DSB sites, they do not play a major role in NHEJ-mediated repair. Consistent with this, analysis of NHEJ events at the non-transcribed Chr3_92 locus, which does not produce diRNAs, also showed no differences between wild-type, *dcl4* and *ago2* mutant plants (Supplementary Fig. S3).

**Figure 4.**
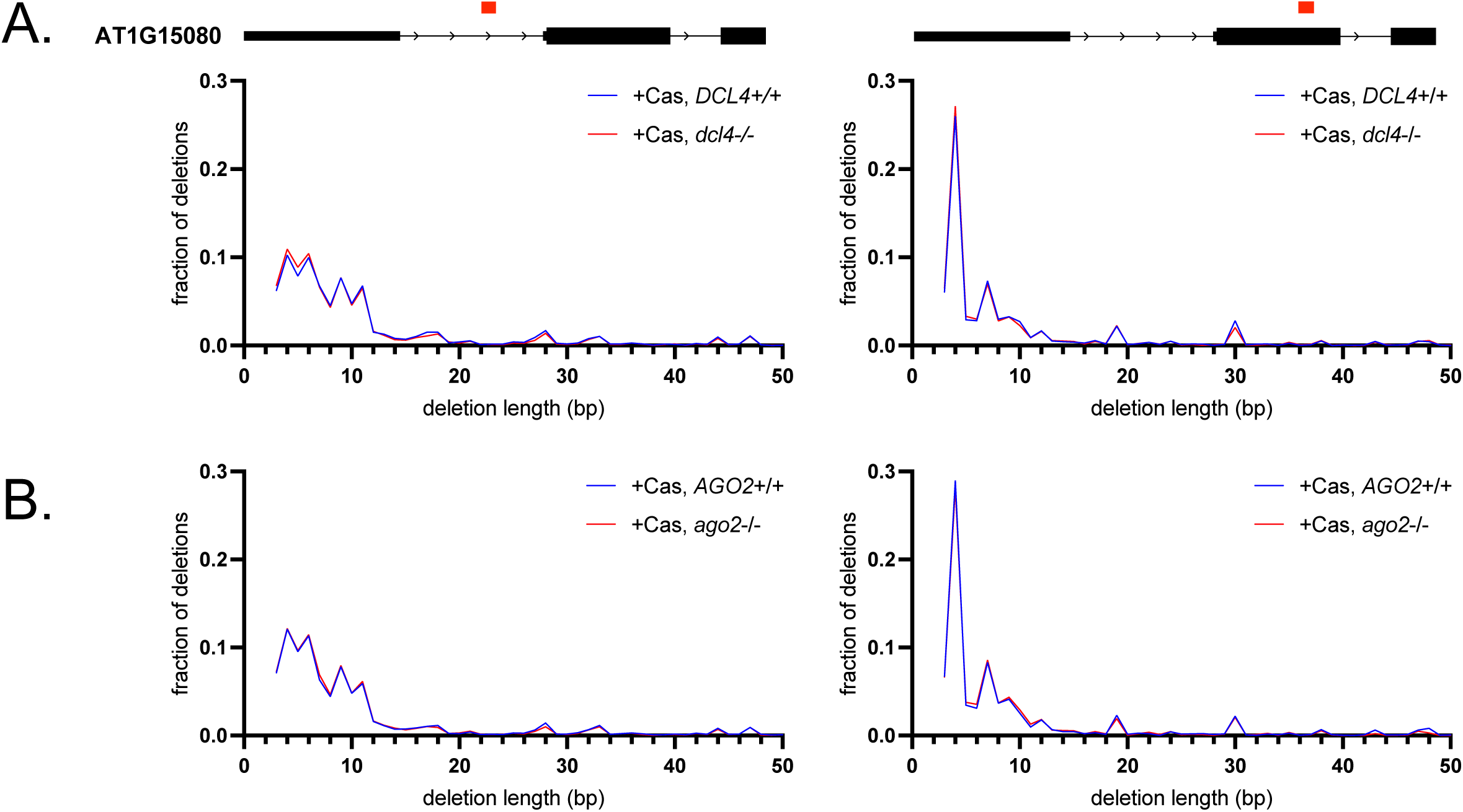
diRNAs are not involved in Non-Homologous End Joining. Frequency distributions of deletion lengths at two distinct cut-sites on AT1G15080 (*LPP2*) in wild-type, (A) *DCL4* or (B) *AGO2* mutant backgrounds. End Joining repair patterns were obtained from a subset of 50k reads. Graphs show reads with a deletion >2 bp as smaller deletions could not be differentiated from sequencing errors.

To further investigate whether diRNAs affect alternative end-joining pathways, we analyzed microhomology-mediated end joining (MMEJ) at the same DSB sites. For each deletion event, we compared the sequences immediately upstream and downstream of the deleted region to identify stretches of microhomology, allowing us to determine the length of microhomology used during repair. We then compared microhomology length distributions between wild-type, *dcl4* and *ago2* mutant plants (Fig. 5 and Supplementary Fig. S4). Remarkably, microhomology usage profiles were highly similar between the three genotypes across all target sites, indicating that diRNAs do not play a major role in MMEJ-mediated repair at endogenous DSBs (Fig. 5).

**Figure 5.**
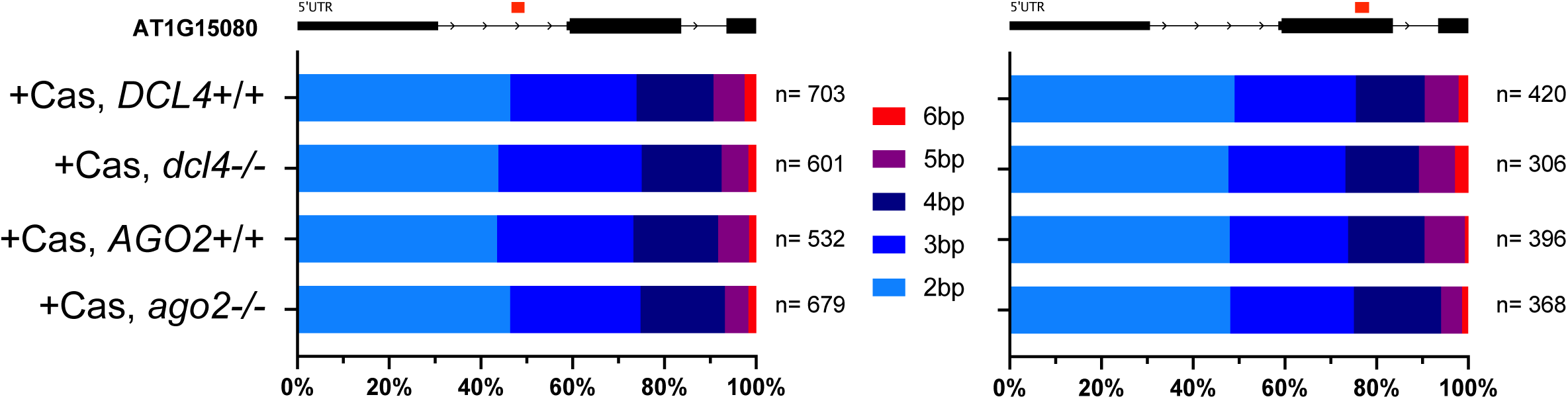
diRNAs have little to no impact on Microhomology-Mediated End Joining. Proportion of reads with a deletion associated with the use of sequence microhomology, at each CRISPR/Cas12 cut-sites of the expressed gene AT1G15080 (*LPP2*), in wild-type, *DCL4* or *AGO2* mutant backgrounds. Size of sequence microhomology was evaluated by assessing sequence motifs present in both the deleted portion of read and directly at the vicinity of the DNA double-strand break in an initial subset of 50k reads, n indicates the total number of reads with a microhomology between 2 and 6 bp.

While our data indicate that the modes of repair for NHEJ and MMEJ are not altered, they do not directly address whether these pathways are downregulated or upregulated. Indeed, our approach rather focuses on the repair profile but does not allow us to quantitatively assess the efficiency of NHEJ and MMEJ. To address this limitation, we estimated repair efficiency by calculating the proportion of unmodified and modified reads in our sequencing data. We reasoned that defects in NHEJ and/or MMEJ would likely affect the frequencies of these reads. This analysis was performed for both wild-type plants and *ago2* mutants, in which diRNAs are produced but not bound, at our four target loci (eight cut sites in total). Notably, the frequencies of unmodified and modified reads were highly similar between wild-type plants and *ago2* mutants across all eight cut sites, indicating that diRNAs do not affect end-joining repair. As expected, however, these frequencies varied significantly between cut sites, clearly indicating that cutting efficiency differs between genomic regions (Supplementary Fig. S5).

Taken together, our data indicate that loss of diRNA production or AGO2-mediated diRNA function has little to no detectable impact on the outcome or apparent efficiency of end-joining repair, including both NHEJ and MMEJ, at the DSB sites analyzed. Thus, despite their robust production at euchromatic DSBs, diRNAs do not appear to play a major role in shaping end-joining repair outcomes at these endogenous genomic loci.

### Endogenous diRNAs contribute to RAD51 recruitment to DSBs

Our data above suggest that diRNAs play no major role in NHEJ and MMEJ repair. However, previous studies in plants and mammals have suggested that diRNAs may promote HR repair through AGO2 by facilitating the recruitment of RAD51 to the site of the DNA break (18, 34). To investigate the genome-wide contribution of AGO2 to HR repair, we examined RAD51 focus formation following treatment with Mitomycin C (MMC). MMC induces DNA interstrand crosslinks that can lead to DNA breaks requiring homologous recombination for repair. Consistent with this, HR-defective mutants in *Arabidopsis* are hypersensitive to MMC, and MMC treatment induces numerous RAD51 foci (38–42). We performed RAD51 immunofluorescence staining in root tip nuclei of 5-day-old seedlings treated or not with 30 µM MMC for 6 hours (Fig. 6A). As expected, no or very few foci were detected in root tip nuclei of untreated seedlings (-MMC, Fig. 6A). In contrast, numerous RAD51 foci were detected in nuclei following MMC treatment (+MMC). Remarkably, no difference was observed between wild-type plants and *ago2* mutants: 31% (n = 78/248 cells) and 34% (n = 128/374 cells) of nuclei were positive for RAD51, with an average of 6.3 and 6.8 RAD51 foci per cell, respectively (Fig. 6A). Together, these results indicate that AGO2 is not required for RAD51 focus formation in response to MMC-induced DNA damage, suggesting that AGO2 does not have a major genome-wide role in HR repair under these conditions. However, MMC induces DNA damage throughout the genome and is not expected to specifically generate 21-nt diRNAs at the resulting lesions. We therefore next asked whether AGO2 contributes to HR repair in a diRNA-dependent context by examining HR at the *LPP2* locus which produces diRNAs upon DNA damage. To address this question, we adapted the experimental design used above for NHEJ and MMEJ to specifically monitor HR events during the DSB break–repair cycle. In the experiments examining end-joining repair, all analyses were performed using 10-day-old plants cultivated at 29 °C, a condition under which the Cas12a enzyme is constitutively active, leading to a continuous cycle of DSB induction and repair. This cycle persists as long as DSBs are accurately repaired, either by NHEJ or by HR using the sister chromatid during the G2 phase. Exit from this cycle occurs upon imperfect repair, which disrupts gRNA target recognition and consequently prevents further Cas12a-mediated DSB induction. As a result, HR cannot be quantitatively assessed in this experimental set-up, as the final molecular outcome primarily reflects imperfect NHEJ events. To specifically investigate homologous recombination, we aimed to analyze repair events during the break–repair cycle rather than at its endpoint. The Cas12a system offers temporal control of DSB induction, as its enzymatic activity is enhanced at 29 °C (43, 44). Accordingly, 10-day-old plants grown at 23 °C were shifted to 29 °C for 4 hours to transiently induce DSB formation. A subset of these breaks is expected to occur during the G2 phase, thereby falling within a temporal window compatible with repair by RAD51-mediated homologous recombination using the sister chromatid. To directly assess the role of endogenous diRNAs in homologous recombination, we used RAD51 focus formation, detected by immunofluorescence, as a proxy for HR repair. We compared wild-type plants and *ago2* mutants using Cas12a lines targeting the *LPP2* locus. As expected, no or very few RAD51 foci were detected in the absence of Cas12a in either genotype (-Cas, AGO2+/+ and -Cas, *ago2−/−*, respectively; Fig. 6B). In contrast, RAD51-positive nuclei were readily detected in both Cas12a-containing plants (+Cas, AGO2+/+ and *+*Cas*, ago2−/−;* Fig. 6B) confirming that the absence of AGO2 does not prevent RAD51 focus formation at DSB sites. Most RAD51-positive nuclei contained a single focus, rarely two, consistent with the induction of a targeted DSB at a single locus. To detect more subtle differences between the genotypes, we quantified RAD51 focus formation as the number of foci per 100 nuclei. Following Cas12a activation, we detected an average of 1.9 RAD51 foci per 100 nuclei in wild-type plants (Fig. 6B). This number was reduced by 42% in *ago2* mutants, with 1.1 foci per 100 nuclei, representing a modest but statistically significant decrease (Fig. 6B). These results are consistent with a role for AGO2 in promoting RAD51 recruitment to DSBs.

**Figure 6.**
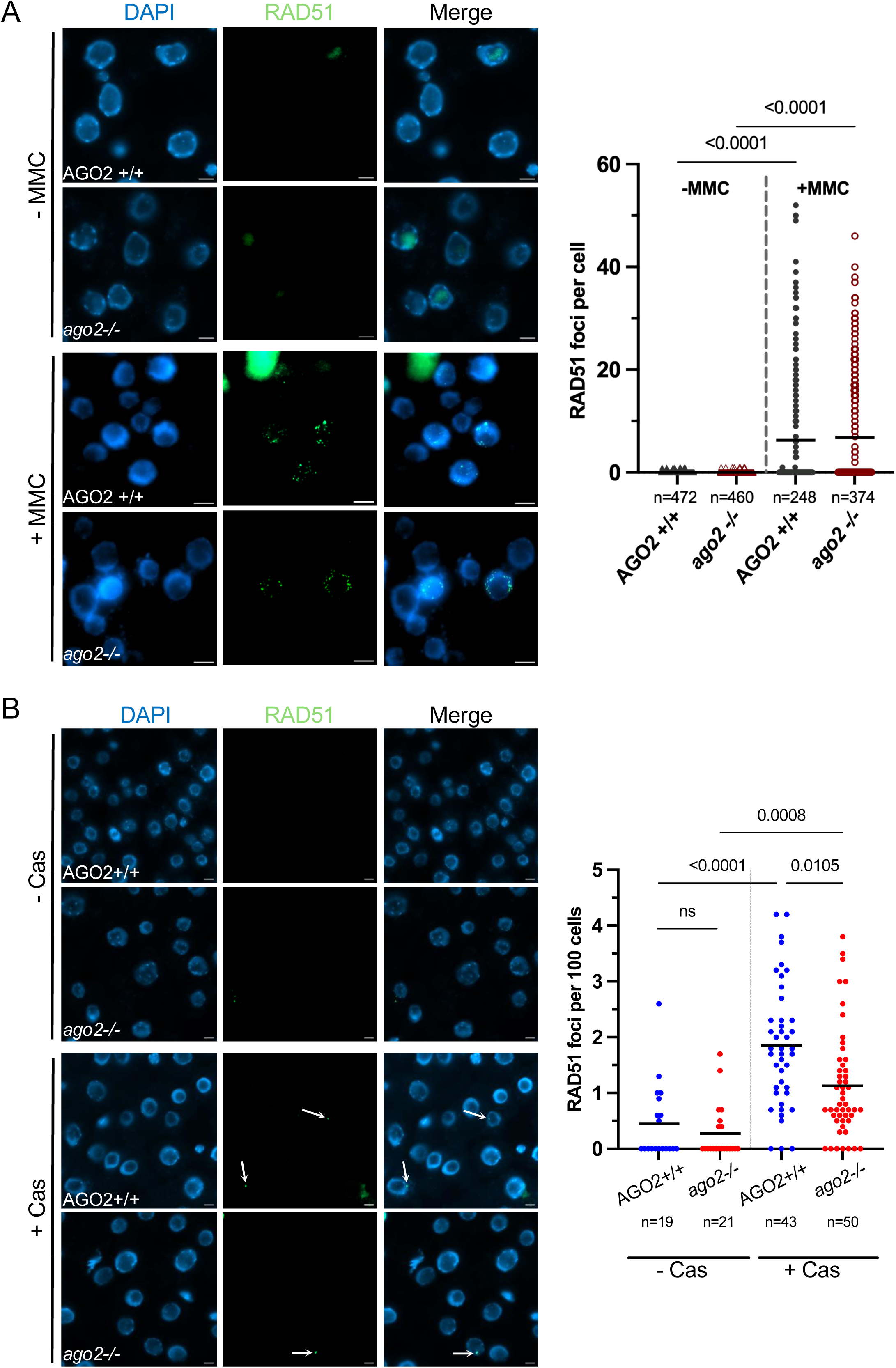
Endogenous diRNAs contribute to RAD51 recruitment to DSBs. (A) Representative images of RAD51 immunolocalization in root tip nuclei of five-day-old wild-type (AGO2 +/+) and *ago2-1* mutant (*ago2 −/−*) seedlings treated with 30 µM mitomycin C for 6 hours (+MMC) or left untreated (−MMC). The corresponding quantification of RAD51 foci per cell is shown in the graph for each genotype and treatment condition. Each dot represents one individual cell, and mean is represented by black line. Statistical analysis was performed using Kruskal Wallis test (two-sided). (B) Immunolocalization of RAD51 in root tip nuclei of 10-day-old wild-type (AGO2 +/+) and *ago2-1* mutant (*ago2 −/−*) seedlings. Quantification of RAD51 foci per 100 cells for each genotype is shown in the graph. Each dot represents the number of RAD51 foci per 100 cells following CRISPR/Cas12a-induced DSBs at the AT1G15080 (*LPP2*) locus (+Cas), in a WT or *AGO2* mutant background. Mean is represented by a black line. Statistical analysis was performed using Kruskal Wallis test (two-sided), ns indicates no significant difference. Scale bar: 5 µm.

Collectively, these results suggest that AGO2 contributes to RAD51 recruitment at Cas12a-induced DSBs within a coding region, but is not generally required for RAD51 focus formation following MMC treatment. Given the random distribution of MMC-induced DNA damage, most resulting lesions would be expected to occur outside coding regions and therefore are unlikely to generate 21-nt diRNAs, the differential effect of AGO2 deficiency between the two experimental conditions is consistent with a contribution of endogenous diRNAs to RAD51 recruitment at DSBs.

## Discussion

The presence of diRNAs at endogenous loci remains subject to debate. To date, only one study in *Drosophila* has reported their accumulation following a DSB at an endogenous, unique gene locus (45). Their biological function also remains unclear. In this study, we investigated whether 21-nt diRNAs are produced at endogenous DSBs in *Arabidopsis thaliana* and whether they contribute to DSB repair. Using CRISPR/Cas12a, we generated DSBs at loci with distinct transcriptional contexts and examined both diRNA production and DSB repair outcomes. We found that 21-nt diRNAs are produced at actively transcribed loci but not at an untranscribed locus, indicating that transcription is required for their biogenesis. Their production was dependent on DCL4. Long-read DNA sequencing revealed no detectable effect of diRNAs on DSB repair by either end joining or microhomology-mediated end joining, but a modest effect on homologous recombination. Moreover, diRNA production was associated with reduced expression of damaged genes, consistent with a role in post-transcriptional gene silencing. Collectively, these findings support a model (Fig. 7) in which diRNAs contribute to the post-transcriptional silencing of damaged genes, potentially limiting the expression of broken loci until repair is completed (Fig. 7). In parallel, as previously suggested and further supported by our data, diRNA-associated AGO2 may favor RAD51 recruitment (Fig. 7), thereby promoting the use of a less mutagenic repair pathway when a transcriptionally active gene is damaged.

**Figure 7.**
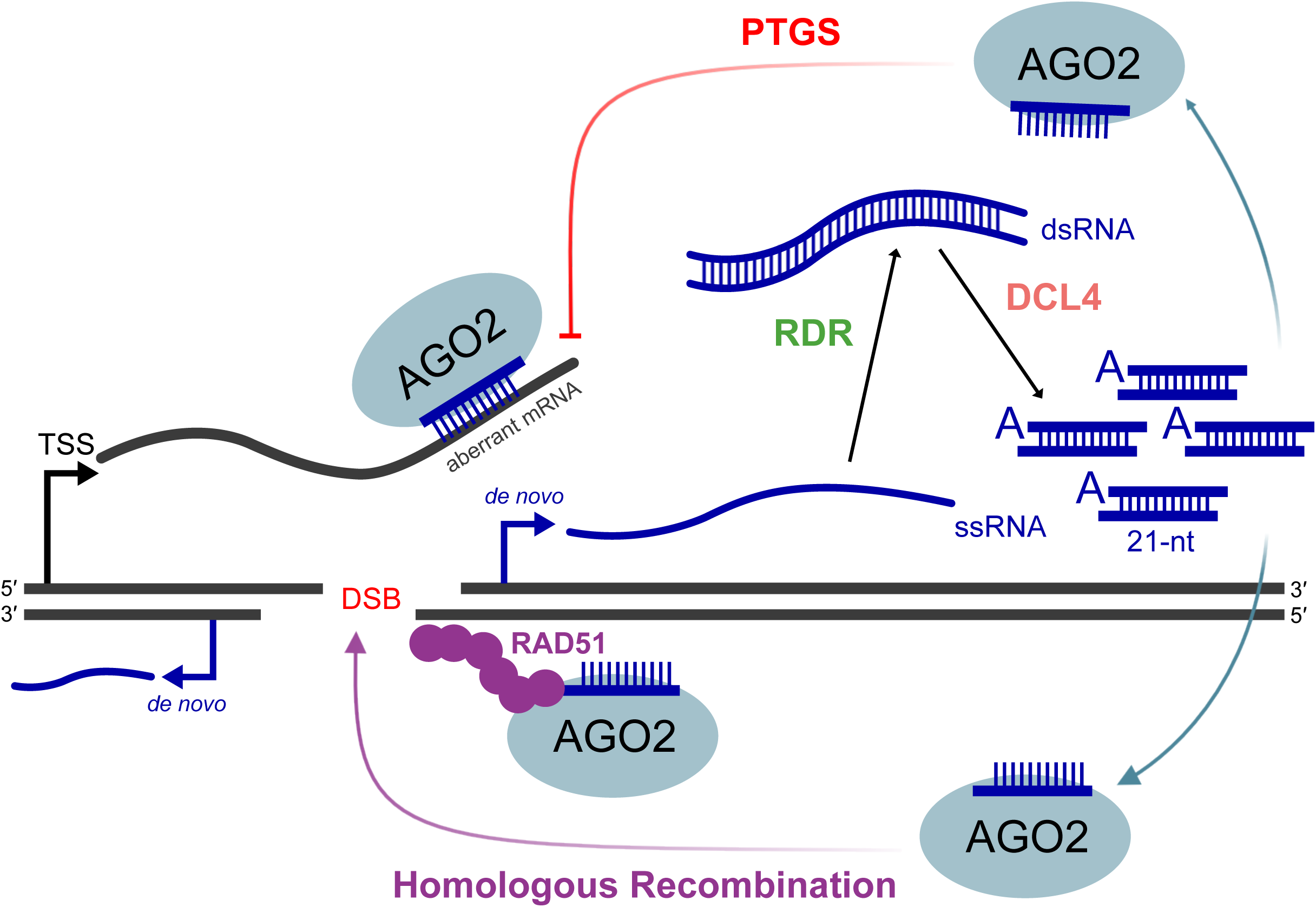
Model depicting the roles of endogenous diRNAs in DSB repair and PTGS of damaged genes. We propose a model that refines and extends previous models of the roles of endogenous diRNAs in DSB repair. Our data show that endogenous 21-nt diRNAs are produced at endogenous sites of active transcription in a DCL4-dependent manner. Consistent with previous studies, we propose that single-stranded RNAs (ssRNA) produced at regions flanking the DSB are subsequently converted into a double-stranded RNA precursor (dsRNA) by RNA-dependent RNA polymerases (RDR), which is subsequently processed by DCL4 into 21-nt diRNAs. These diRNAs are then loaded into AGO2. Our data further support this previously proposed pathway, as indicated by the presence of a 5′ adenosine (5′ A) in these diRNAs, a characteristic associated with AGO2 loading. We propose that the resulting diRNA-AGO2 complexes contribute to post-transcriptional gene silencing (PTGS) of their originating damaged gene, thereby limiting aberrant mRNA accumulation. Furthermore, building on previous evidence that diRNA-AGO2 complexes favor the local recruitment of RAD51 to DSBs, our data provide further support for a role of diRNAs in promoting repair by Homologous Recombination. Collectively, our results both support the previously proposed role of diRNAs in DSB repair and extend the model by implicating endogenous diRNAs in the PTGS of damaged genes. We note that the interaction of the diRNA-AGO2 complex at regions flanking the DSB may require an RNA transcript, the identification of which remain to be determined.

### diRNAs are produced at endogenous transcribed loci in a DSB-dependent manner

Accumulating evidence has revealed an unexpected role for RNA molecules, and particularly small RNAs, in DNA repair across several organisms, including *Neurospora crassa*, *Arabidopsis thaliana*, *Drosophila* and mammalian cells (6–8, 15). However, 21-nt diRNAs have mostly been detected at transgenic or repetitive ribosomal DNA sequences, whereas their production at unique endogenous genic or intergenic loci has been more difficult to demonstrate (6, 15, 46). In *Arabidopsis*, 21-nt diRNAs were initially identified from sequences flanking a site-specific DSB introduced into a transgene (15). However, subsequent analyses failed to detect diRNAs at eight endogenous target genes (including two containing repeat sequences) in *Arabidopsis* and five endogenous target genes in rice, with diRNAs detected only at a highly transcribed transgenic locus (19). These findings led to the proposal that high levels of RNA polymerase II transcription might be required for efficient diRNA production and that components of the post-transcriptional gene silencing (PTGS) pathway, including DCL4, are involved in their biogenesis.

More recently, sense and antisense transcripts were detected around DSBs induced by the restriction enzyme SbfI in approximately 30% of the *Arabidopsis* genes analyzed, with their presence strongly correlated with gene expression levels (20). Although these damage-induced transcripts were frequently bidirectional and dependent on RNA polymerase II, they were identified by poly(A)+ RNA sequencing and therefore correspond to long RNA transcripts rather than the 21-nt diRNAs originally described. This distinction is important because the production of long DSB-induced transcripts are RNA polymerase II-dependent, whereas small diRNAs production has been linked to RNA polymerases IV and V. Consistent with this, the presence of long DSB-induced transcripts does not necessarily result in the production of 21-nt diRNAs (46).

Our results provide, to our knowledge, the first evidence that 21-nt diRNAs can be produced at endogenous loci in *Arabidopsis* (Fig. 1). We detected both sense and antisense 21-nt diRNAs at six DSB sites located within three actively transcribed genes, whereas no diRNAs were detected at two DSB sites within a non-transcribed locus. These results indicate that endogenous, transcriptionally active loci can produce 21-nt diRNAs in response to a DSB and support a role for transcriptional activity in diRNA biogenesis.

So, why do we detect diRNAs at DSBs at endogenous loci, whereas previous studies did not? There is clearly a link with transcription, since we only detected diRNAs at transcribed loci. However, transcription might be necessary but not sufficient, as the effect does not appear to be simply determined by transcriptional activity. For example, Miki et al. also targeted endogenous genes but did not detect diRNAs (19). Similarly, long RNAs were detected at some DSBs within genes, but not at all genes, only around 30% of the loci analyzed (20). Although the latter study detected long RNAs rather than diRNAs, diRNAs are thought to be generated through processing of these long RNA precursors, and therefore would not be expected at loci where long RNAs are not produced. We therefore favor the hypothesis that high transcriptional activity, potentially in combination with specific chromatin features, is required for efficient diRNA production (see below). This is supported by the detection of diRNAs at transgenes and repetitive ribosomal DNA sequences, which are generally highly transcribed (19, 46).

We also do not think that the type of DNA-cutting system is the main determinant. A variety of systems have been used to induce DSBs, including the I-SceI meganuclease, restriction enzymes, TALENs, Cas9 and, in our study, Cas12a. Cas12a differs from Cas9 and TALENs in that it generates staggered, cohesive DNA ends, whereas Cas9 and TALENs generally produce blunt ends. However, this difference is unlikely to explain diRNA production, since diRNAs have been observed at the GU-US transgene following both Cas9- and I-SceI-induced DSBs (19), and I-SceI also generates cohesive ends. Thus, the nature of the DNA-break extremities does not appear to be a major determinant. Consistent with this, 21-nt damage-induced siRNAs have also been detected after UV-induced DNA lesions (21), suggesting that their production is not restricted to a particular type of DNA break. Another possibility is that the higher cutting efficiency of Cas12a at 29°C contributes to the phenotype. However, this alone is unlikely to explain our results. In Miki et al., diRNAs were detected at the GU-US transgene following Cas9-induced DSBs, but not at their endogenous targets, despite likely comparable cutting efficiencies (19). Therefore, cutting efficiency alone cannot account for the difference.

Overall, we favor a combination of factors rather than a single determinant. In our system, efficient Cas12a cutting, together with high transcriptional activity and a favorable chromatin environment, may promote diRNA production. The relatively high growth temperature of 29°C may further contribute by affecting transcription and/or DSB formation. This combination of factors could explain why we detect diRNAs at endogenous DSBs whereas previous studies did not.

### Origin and biogenesis of the 21-nt diRNAs

The comparable abundance of sense and antisense diRNAs observed at each of the three loci suggests that diRNAs may originate from double-stranded RNA (dsRNA) precursors. However, the identity of these precursors remains unclear. The absence of diRNAs when DSBs were introduced in a non-coding region (Chr3_92 target), together with their production at genic loci, points to transcription as an important determinant of diRNA biogenesis. Consistent with this, a study using a transgene showed that RNAPII transcription of the damaged sequence was required for efficient diRNA production, suggesting that a transcript generated from the damaged locus could serve as a substrate for dsRNA synthesis (19). However, the detection of diRNAs both upstream and downstream of the DSB in our three genes cannot be readily explained by processing of a single pre-existing transcript initiated at the transcription start site. One possibility is that DSBs promote local transcription initiation, as reported in *Arabidopsis*, where damage-induced RNAs were frequently initiated at DSB sites within actively transcribed genes and detected in both directions around the break (20, 46–50). Nevertheless, DSB-associated transcription remains debated, as other studies have failed to detect RNAPII recruitment at DSBs (51, 52), indicating that the contribution of *de novo* transcription to diRNA production remains unresolved. Interestingly, our observation that diRNAs are restricted to the exon or intron containing the DSB further suggests that RNA processing may influence the availability of diRNA precursors. In *Drosophila*, splicing factors were shown to be required for efficient siRNA production at DSBs, with DSB-induced defects in co-transcriptional splicing proposed to promote the accumulation of RNA substrates for small-RNA biogenesis (45). Finally, the dependence of diRNA production on DCLs (Fig. 2) and RDRs, together with the reported involvement of Pol IV, supports the idea that transcripts generated or retained at the damaged locus may subsequently enter established plant small-RNA pathways (15, 19). Further work will be required to determine whether these precursors correspond to pre-existing transcripts, damage-induced RNAs, or unprocessed RNA species generated at the DSB.

### 21-nt diRNAs do not play a major role in NHEJ or MMEJ

To date, evidence for a role of DSB-induced small RNAs in NHEJ remains largely indirect. One study reported that impairment of the small RNA pathway affected the efficiency of NHEJ (53). However, this conclusion was based on the use of *Pol IV* and *dcl3* mutants, which are components of the small RNA biogenesis pathway. These mutants have pleiotropic effects, meaning that they impact not only 21-nt small RNAs but also other processes. Thus, this does not directly demonstrate that siRNAs themselves act at the DNA break or interact with NHEJ factors. Here, we directly investigated the contribution of DSB-induced small RNAs by using *dcl4* and *ago2* mutants, which impair diRNA production and disrupt the function of their putative effector, respectively. We did not observe any clear effect of their loss on NHEJ or MMEJ pathway usage or on the resulting repair outcomes (Fig. 4 and 5). To our knowledge, this is the first direct assessment of the contribution of diRNAs to NHEJ and MMEJ. Our results indicate that they do not play a major role in pathway usage or repair outcomes under the conditions tested.

### 21-nt diRNAs potentially promote HR at gene-associated DSBs

If AGO2 plays a major role in homologous recombination (HR), its loss would be expected to compromise the cellular response to DNA damage. Indeed, the expression of *Arabidopsis thaliana* AGO2 is strongly induced under genotoxic stresses such as γ-irradiation and zeocin treatment (54, 55). However, evidences from *Arabidopsis thaliana* are inconsistent. While some studies have reported increased sensitivity of *ago2* mutants to DNA-damaging agents such as mitomycin C, other have observed increased resistance to zeocin, with the phenotype depending on the mutant allele analyzed, highlighting the complexity of AGO2-associated DNA damage responses (56, 57). Nevertheless, several lines of evidence support a role for AGO2 in DSB repair. In *Arabidopsis*, AGO2 immunoprecipitation experiments demonstrated that diRNAs are loaded onto AGO2, with a strong reduction of diRNAs in *ago2* mutants (15), whereas in *Drosophila*, the small-RNA response induced by DSBs is impaired in *ago2* mutant cells (58). More directly, reporter-based assays have shown that depletion of AGO2 reduces HR efficiency in both plants and mammalian cells (15, 34, 35). In mammals, studies using reporter systems have proposed that diRNA biogenesis and AGO2 activity promote HR by facilitating the recruitment of RAD51 and BRCA1 to DSBs and by contributing to local chromatin remodeling through the recruitment of MMSET and TIP60 (34, 35, 59). Consistent with this model, AGO2 has also been implicated in RAD51 localization at DSBs in *Arabidopsis thaliana*, where the diRNA-AGO2 pathway was proposed to facilitate IDN2 recruitment and the replacement of RPA by RAD51 (18), and to contribute to the recruitment of the SMC5/6 complex at damaged sites (16). However, the contribution of diRNAs and AGO2 to DSB repair is not universal across organisms or repair contexts. For example, depletion of Dcr-2 and Ago2 in *Drosophila* did not impair HR, and increasing diRNA production in mammalian cells has even been reported to favor 53BP1 recruitment and NHEJ rather than HR (58, 60). These discrepancies may reflect differences in experimental systems, DSB induction methods, repair substrates, or species-specific functions of small RNAs.

Importantly, our results provide evidence for a role of AGO2 in HR at an endogenous genomic locus rather than in a reporter system (Fig. 6). We observed a 42% reduction in RAD51 accumulation at endogenous DSBs associated with diRNA production in *ago2* mutants (Fig. 6). This reduction mirrors the 44% HR defects observed using the *Arabidopsis* GUS reporter system (15). Thus, our data extend previous reporter-based observations by suggesting that AGO2 may be involved RAD51 recruitment at an endogenous DSB, and that this pathway is not restricted to engineered reporter substrates. Nevertheless, whether the observed reduction is due specifically to the loss of AGO2 itself or indirectly to the absence of AGO2-associated diRNAs cannot be distinguished from our experiments. Further separation of AGO2 functions from diRNA-dependent effects will therefore be required to determine whether AGO2 acts primarily as a diRNA-guided scaffold at DSBs or also contributes to HR through diRNA-independent mechanisms.

Collectively, our findings, together with previous studies, support a model in which 21-nt diRNAs and AGO2 may contribute to HR at endogenous DSBs, at least in part by promoting RAD51 accumulation at sites of DNA damage.

### 21-nt diRNAs contribute to post-transcriptional regulation of transcript accumulation at DSBs

Our results suggest that DSB-induced 21-nt diRNAs negatively affect transcript accumulation at the three endogenous loci analyzed in this study (Fig. 3). Following DSB induction, transcript levels decreased in wild-type plants, whereas this reduction was restored in the *dcl4* mutant, supporting a model in which DCL4-dependent 21-nt diRNAs contribute to the reduction in transcript abundance downstream of DSB formation. These diRNAs also associate with AGO2 (Fig. 2), suggesting that they may act through an AGO2-dependent silencing mechanism. AGO2 is a well-established effector of small-RNA-mediated silencing in plants, and DCL4-dependent 21-nt siRNAs can promote the sequence-specific repression or degradation of homologous transcripts (13, 61–64). Thus, DSB-induced 21-nt diRNAs may provide a link between DNA damage and post-transcriptional regulation of transcripts from damaged loci, potentially through AGO2. In this model (Fig. 7), diRNAs generated in response to a DSB would be loaded into AGO2 and guide the recognition and subsequent repression or degradation of transcripts originating from the damaged locus, thereby providing a direct mechanism linking DNA damage to post-transcriptional regulation of gene expression.

However, we cannot yet conclude that the observed decrease in transcript abundance results from canonical PTGS. RT-qPCR measurements of steady-state RNA cannot distinguish increased RNA degradation from reduced transcription, altered RNA processing or other DSB-associated effects on gene expression. This distinction is particularly relevant because DSBs can themselves induce local transcriptional repression, including reduced RNAPII occupancy and transcriptional output (14). Moreover, whether AGO2-bound 21-nt diRNAs can influence chromatin organization or transcription at the damaged locus remains unknown. Further experiments measuring nascent transcription, RNA stability and AGO2 dependence will therefore be required to establish whether the decrease in transcript accumulation is indeed mediated by PTGS. Nevertheless, our findings identify a DCL4-dependent, AGO2-associated 21-nt diRNA pathway that accompanies reduced transcript accumulation following DSB formation, suggesting that diRNAs may represent a previously unrecognized mechanism coupling DNA damage to post-transcriptional gene regulation at endogenous loci.

## Acknowledgments

We acknowledge the technical services of the iGReD, in particular the Clermont-Ferrand Imagerie Confocale (CLIC), and Anipath platforms, for their help with microscopy and image analysis, as well as the Bioinformatics Platform (BIM). We are grateful to the Mésocentre Clermont-Auvergne and the plateforme Aubi of the Université Clermont Auvergne for providing help, computing and storage resources. The authors acknowledge the support received from the International Research Center on Sustainable AgroEcosystems (ISITE CAP20-25). We also thank Peter Schlögelhofer for providing the RAD51 antibody and all members of the Meiosis, Recombination and Adaptation group for their help and discussions.

## Author contributions

E.M., C.W., J.V., and S.L. produced data. E.M., C.W., S.L., H.S. and O.D.I. analyzed the data. E.M., C.W., S.L., and O.D.I. conceived and designed the experiments. E.M., S.L., and O.D.I. wrote the manuscript with input from all authors.

## Supplementary data

Supplementary data are available at NAR online

## Data availability

All data underlying this article are available in the article and its online Supplementary data. The sequence files are available under BioProject accession number PRJNA1541105 at the NCBI Sequence Read Archive (SRA). NHEJ and MMEJ analysis were performed using an in-house Python and R script publicly available at https://github.com/chrlesw/dsbrepair. Additional datasets generated during this study are available from the corresponding authors on request.

---

**Supp Fig S1. RNAseq suggests PTGS at damaged genes upon diRNA production** mRNA abundance of CRISPR/Cas12a-targeted genes was assessed by RNA sequencing in 10-day-old plants producing diRNAs (*DCL4*+/+) or not (*dcl4*−/−). (A) Data were normalized as Transcripts Per Million (TPM). The “+DSB” annotation refers to lines expressing a functional CRISPR/Cas12 system (+Cas) targeting the evaluated gene, while “-DSB” is related to uninduced lines producing diRNAs. (B) RNAseq coverage tracks for the CRISPR/Cas12a-targeted genes AT1G15080, AT5G41150, and AT3G05210 in 10-day-old plants. Screenshot of the Integrative Genomics Viewer (IGV) genome browser showing coverage for uninduced lines (-DSB, gray), lines expressing a functional CRISPR/Cas12a and producing diRNAs (+DSB, DCL4+/+, blue), and lines lacking DCL4 and thus not producing diRNAs (+DSB, *dcl4−/−,* red). Gene models are represented below each track, with exons represented as black boxes and introns as connecting lines. The y-axis indicates coverage in Counts Per Million (CPM).

**Supp Fig S2. End-Joining repair patterns at the AT5G41150 and AT3G05210 target loci** End Joining repair patterns of CRISPR/Cas12-targeted expressed genes (A) AT5G41150 and (B) AT3G05210. Frequency distributions of deletion lengths at two distinct cut-sites were represented from wild-type, *DCL4 (dcl4−/−)* or *AGO2 (ago2−/−)* mutant plants. End Joining repair patterns were obtained from a subset of 50k reads.

**Supp Fig S3. End-joining repair patterns at the Chr3_92 locus** Frequency distributions of deletion lengths at two distinct cut-sites on the pericentromeric repeat-rich target “Chr3_92” in a wild-type, (A) *DCL4* or (B) *AGO2* mutant backgrounds. End Joining repair patterns were obtained from a subset of 50k reads.

**Supp Fig S4. MMEJ analysis at the AT5G41150, AT3G05210 and Chr3_92 target loci** Proportion of reads with a deletion associated with the use of sequence microhomology, at two distinct CRISPR/Cas12 cut-sites on (A) AT5G41150, (B) AT3G05210 and (C) the pericentromeric repeat-rich target “Chr3_92”, in wild-type, *DCL4* or *AGO2* mutant backgrounds; n indicates the total number of reads with a microhomology between 2 and 6 bp from an initial subset of 50k reads.

**Supp Fig S5. CRISPR/Cas12a repair efficiency in wild-type and *ago2* mutants** CRISPR/Cas12a cutting efficiency at each cut-site (CS) was estimated by representing the proportion of unmodified reads and reads bearing a deletion strictly superior to 2 bp, resulting from EJ error-prone repair. Subsets of 50k reads for each target (AT1G15080, AT5G41150, AT3G05210) were examined.

**Supplementary Table S1.**
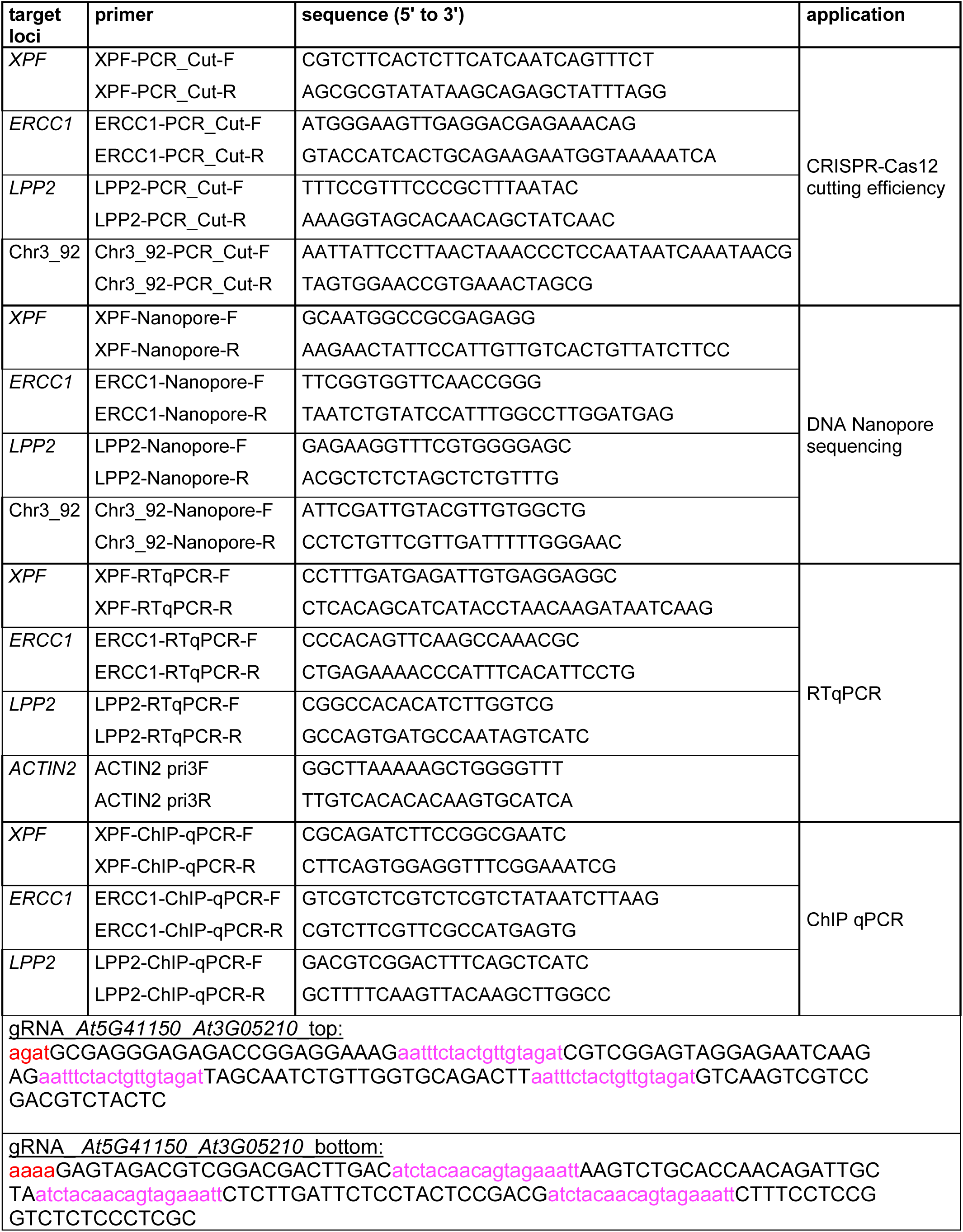
Sequences of the gRNAs and primers used in this study. crRNA sequences are in upper-case, fnCas12a direct repeats are magenta, BbsI restriction sites are red

